# *mirror* determines the far posterior domain in butterfly wings

**DOI:** 10.1101/2024.02.15.580576

**Authors:** Martik Chatterjee, Xin Y. Yu, Noah K. Brady, Gabriel C. Hatto, Connor A. Amendola, Robert D. Reed

## Abstract

Insect wings, a key innovation that contributed to the explosive diversification of insects, are recognized for their remarkable variation and many splendid adaptations. Classical morphological work subdivides insect wings into several distinct domains along the antero-posterior (AP) axis, each of which can evolve relatively independently to produce the myriad forms we see in nature. Important insights into AP subdivision of insect wings comes from work in *Drosophila melanogaster*, however they do not fully explain the diversity of AP domains observed across broad winged insects. Here we show that the transcription factor *mirror* acts as a selector gene to differentiate a far posterior domain in the butterfly wing, classically defined as the vannus, and has effects on wing shape, scale morphology, and color pattern. Our results support models of how selector genes may facilitate evolutionarily individuation of distinct AP domains in insect wings outside of *Drosophila*, and suggest that the *D. melanogaster* wing blade has been reduced to represent only a portion of the archetypal insect wing.

## RESULTS AND DISCUSSION

Insect wings vary dramatically in their size and morphology. Butterflies and moths in particular have attracted attention for their rapidly evolving wings that show extreme variation in shape and color pattern. Interestingly, the diversity of lepidopteran color patterns appears to follow relatively simple rules, wherein an evolutionarily conserved set spot and stripe pattern elements can vary in size, color, and presence/absence along the antero-posterior (AP) axis.^1,2^ Surveys of wing pattern diversity across butterflies, considering both natural variation and genetic mutants, suggest that wings can be subdivided into five AP domains, bounded by the M1, M3, Cu2, and 2A veins as summarized in **Figure 1A**.^3^ Within each domain there is a strong correlation between pattern variation and wing morphology, and, conversely, between domains there is a relative degree of independence in morphological variation. We know little about how these different AP domains are differentiated during wing development, however.

**Figure 1.**
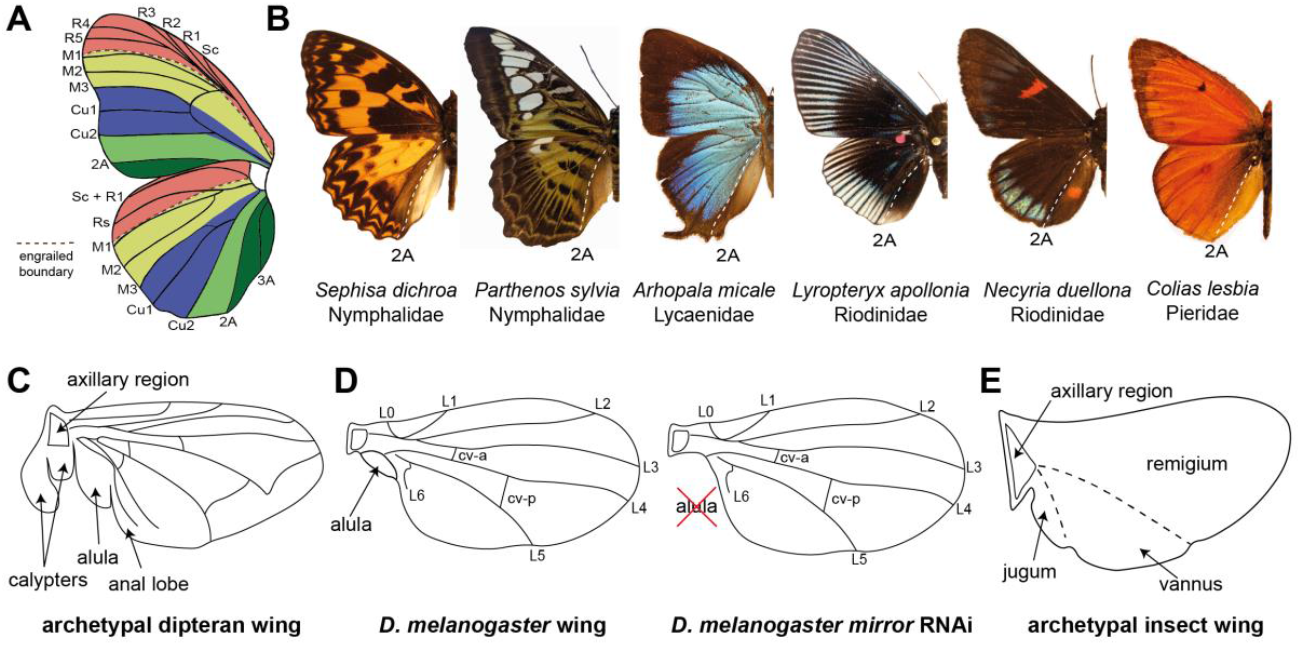
Proposed AP domains of insect wings. (A) Comparative morphology previously identified boundaries associated with color pattern variation, demarcated by the M1, M3, Cu2, and 2A veins.^5^ (B) Examples of the 2A vein marking a clear posterior color pattern boundary across multiple butterfly families. (C) Proximal posterior features of the archetypal dipteran wing include the calypters, which are only found in Calypterata clade, and the alula, a lobe at the base of the wing blade.^21^ (D) *mirror* RNAi knockdowns result in highly specific loss of the alula in *D. melanogaster*.^13^ (E) Snodgrass’ model of the archetypal insect wing specifies three major domains along the anteroposterior axis: the remigium, the vannus, and the jugum. Homologies of these domains with the features of the dipteran wing (C), or butterfly wing pattern boundaries (A), has remained unclear.

Some aspects of the AP wing differentiation process have been extrapolated from *D. melanogaster* wings to butterflies. The transcription factors engrailed and spalt define AP domains in developing fly wings^4-6^ and exhibit expression patterns in butterflies that suggest some functions may be conserved. In butterflies *engrailed* transcription across the entire region posterior to the M1 vein, while its paralog *invected* marks a largely overlapping region between M1 and 2A.^7-9^ *spalt* expression is more dynamic and complex in butterflies, but appears to consistently mark a region between the R2 and M3 veins.^9^ These observations, coupled with the overall size and morphological complexity of butterfly wings, have spurred interest in the question of whether additional AP differentiation mechanisms may underlie AP domain specification in Lepidoptera and other insects.^10,11^ The identification of additional domain-determining transcription factors in broad winged insects would provide a model for the modular diversification of insect wing morphology.

The goal of this study was to identify the molecular basis of AP domain specification in the butterfly wing, posterior of the M1 engrailed boundary. We initially identified the transcription factor *mirror* as a potential candidate selector gene based on previous mRNA-seq studies,^12^ coupled with *D. melanogaster* work showing that *mirror* specifies the alula – a small membranous lobe at the posterior base of the wing in some dipterans (Figure 1C,D).^13,14^ We used *in situ* hybridization to visualize *mirror* mRNA localization in the last-instar imaginal discs of the common buckeye butterfly, *Junonia coenia*, and observed *mirror* expression throughout the wing domain posterior to the 2A vein boundary (**Figures 2A and S1)**. This expression precisely marks the vannus, or anal region **(Figures 1B, E)**, of the butterfly wing – a distinct posterior domain of the wing blade populated by the anal (A) veins and bordered anteriorly by the claval fold between the cubital (Cu) and A veins in the hindwing **(Figure 2B)**. As observed in various representative species across different butterfly families, this region of the hindwing is distinct from the rest of the wing; the color patterns defining the wing blade terminate at the 2A vein, and in many species it is marked by scales that differ in morphology and lack color patterns found in anterior regions of the wing **(Figures 1B and S2A)**. This region has been proposed to represent the posterior-most color pattern domain in butterflies **(Figure 1A, Figure 2B)**.^3^

**Figure 2.**
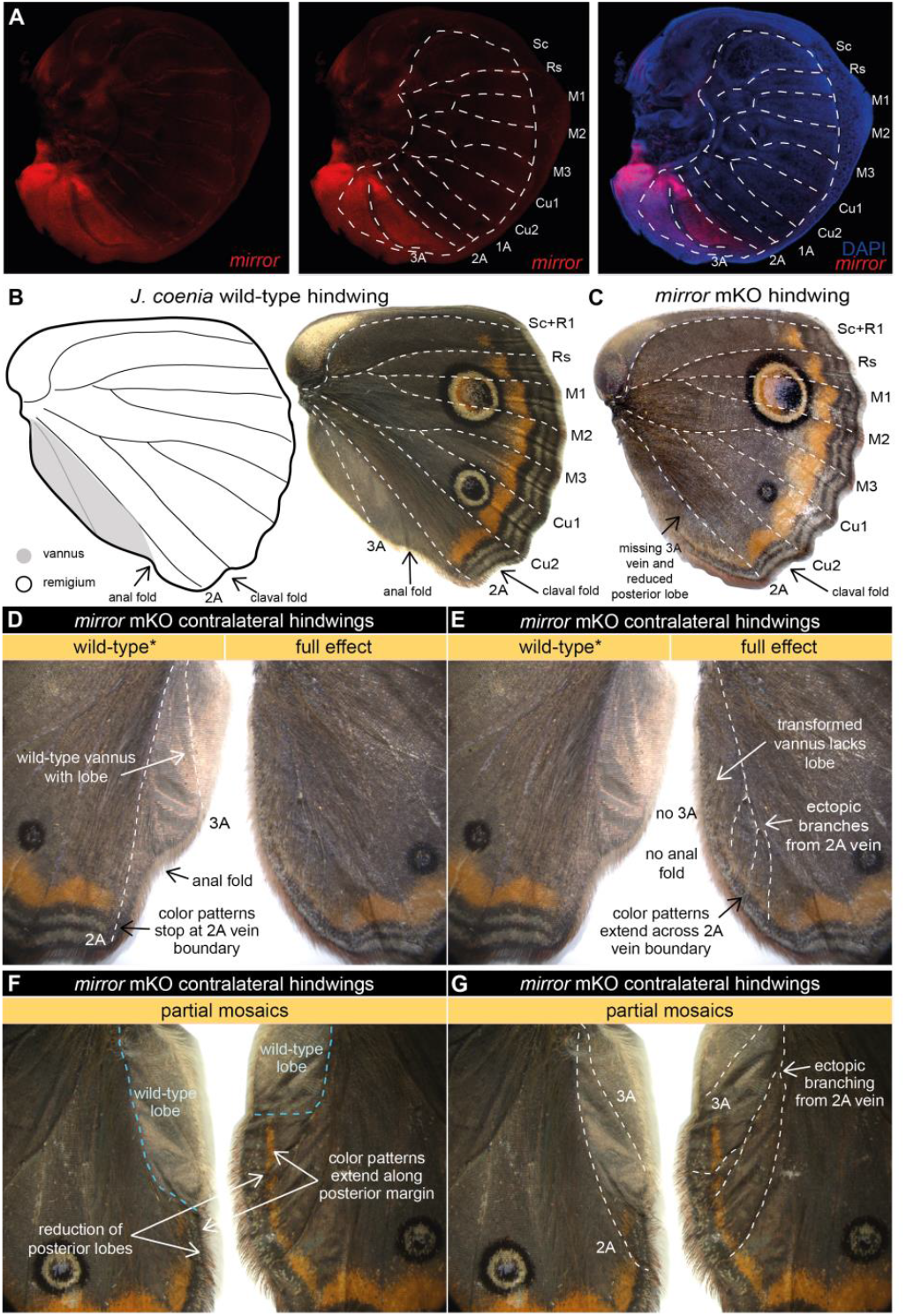
*mirror* determines the identity of the far posterior vannus domain in *J. coenia.* **(A)** In *J. coenia mirror* (red) is expressed posterior to the 2A vein in the late last-instar hindwing imaginal discs, marking the region that will develop into the vannus. See Figure S1 for forewing expression and controls. **(B)** In adult butterfly wings the vannus is the wing field posterior to the 2A vein, shown here in *J. coenia* as a posterior lobe with silvery scales devoid of color patterns found anterior to 2A. **(C)** Targeted mosaic knockouts (mKOs) of *mirror* result in loss of vannus features in *J. coenia*. **(D)** Annotation of wild-type features preserved on one side of a contralateral mKO. **(E)** Annotation of full effect mutant phenotyp in the opposing wing from the individual in (D) highlights loss of lobe, loss of 3A, ectopic branching of 2A, and posterior extension of color patterns. **(F)** Annotation of a partial mKO highlights transformation of posterior lobe and extension of color patterns, both of which terminate at the mutant clonal boundary. **(G)** The individual shown in (F) also shows ectopic branching of 2A in the mutant region, but preservation of 3A in the wild-type region. For additional *mirror* mKO phenotypes see Figures S2-S4.

To functionally assess the role of *mirror* in wing development we generated CRISPR/Cas9 mosaic knockouts (mKOs) targeting the coding region of *J. coenia mirror*. We recovered many mKO mutants that consistently exhibited several distinctive phenotypes **(Table S1)**, but only ever in the *mirror*-expressing posterior vannus region:

1. **Venation:** In mKO mutant hindwings we observed frequent truncation or complete loss of the loss of the 3A vein, as well as discontinuity and ectopic bifurcation of 2A veins **(hindwings: Figures 2C-G, S2, S3; forewings: Figure S4)**.
2. **Wing shape:** mKOs consistently showed shape change along the posterior edge of the wing, especially the hindwing where the projecting lobe characteristic of the vannus was reduced and replaced by a more even, regular wing margin. The anal fold between the 2A and 3A veins, which marks the edge of the vannal lobe, was also lost in mutant clones affecting this wing region **(Figures 2C-G, S2-S3)**.
3. **Color pattern expansion:** In mutant hindwings we consistently observed extension of wing margin color patterns across the 2A vein boundary into the posterior region of the wing. In wild-type wings the submarginal and central symmetry system color patterns terminate at the 2A vein **(dorsal: Figure 2C-G; ventral: Figure S3)**. Remarkably, in *mirror* mKO mutants these entire color pattern systems extend past the 2A vein into the posterior regions of the wing on both dorsal and ventral wing surfaces. Specifically, submarginal bands continue past 2A to follow the posterior wing margin **(Figures 2C-G, S2)**, while central symmetry patterns extend from 2A down to the posterior margin **(Figure S3)**.
4. **Scale morphology:** In most mutants vannus scales lost their distinctive silver structural color and rounded apical edges, and transformed into brown, serrated scales typical of the remigium **(Figure S2A)**. We also observed that scales along mutant wing margins often displayed elongated scales, often manifesting as a dense hair-like matt across the posterior margin **(Figure S2A)**.

The knockout phenotypes we observed illustrate several things about *mirror*’s function in wing development. First, across all mKO replicates we never observed mutant phenotypes anterior to the 2A vein, which suggests that *mirror* effects are limited to the far posterior domain in which we observed *mirror* expression. Second, *mirror* knockouts appear to affect multiple aspects of wing morphology, including scale shape and coloration, overall wing shape, and wing venation. Finally, the loss of *mirror* function resulted in transformation, not loss, of the posterior domain. Even in mKO individuals that appeared to have a mutant phenotype across the entire posterior region, the *mirror*-expressing cells posterior to the 2A vein were not simply lost – the wing field continues well past the 2A vein. Instead, the identities of the post-2A scale cells changed in color and structure. In this respect, the expansion of multiple color pattern systems into the posterior domain of mutants was particularly compelling because the loss of central symmetry system and submarginal patterns in the post-2A vannus region is commonly observed in Lepidoptera **(Figure 1B)**. The ability to reconstitute these complex color pattern elements in the posterior domain by knocking out a single transcription factor leads us to speculate that their patterning information may occur in a latent form that is masked by *mirror* effects. Together, our expression and knockout data suggest that *mirror* acts as a selector gene necessary to define the identity of the far posterior wing domain.

Identifying *mirror* as a vannus-specifying factor is of interest for a few reasons. First, it provides a molecular explanation for how different AP domains of the butterfly wing may be individuated to independently evolve their own shape and color pattern variations. Previous authors have proposed the existence of such individuated domains, and speculated that they may be specified by selector genes.^3,10^ Our data provide experimental support for this model, and now motivate us to identify additional factors that may specify other domain boundaries between the M1 and A2 veins.

Next, the role of *mirror* in specifying the vannus provides an important link between molecular developmental biology and Snodgrass’ classical anatomical designations of the insect wing fields – i.e., the remigium, the vannus, and the jugum **(Figure 1E)**.^15^ The remigium and the vannus are dominant features of most insect wings, while the jugum is smaller lobe-like feature that is reduced or lost in many clades. The remigium is the major anterior wing blade that is directly articulated by thoracic motor musculature to power flight, and is typically divided from the vannus by a distinct wing fold (e.g. the “anal fold” in butterflies). In many insect orders the vannus is a prominent fan-like structure, which can be a greatly enlarged gliding surface some hemimetabolous clades such as Orthoptera (crickets and grasshoppers), Dictyoptera (mantids and roaches), Phasmatodea (stick insects), and Plecoptera (stoneflies). Importantly, *mirror* knockdown in the hemimetaboulous milkweed bug *Oncopeltus fasciatus* causes developmental defects in the claval and anal furrows in the posterior wing,^16^ which leads us to infer that *mirror*’s role in determining the vannus may be deeply conserved in insects. This prompts us to speculate that the developmental individuation of the posterior domain by *mirror* facilitated the remarkable diversification of the vannus across the insects.

Finally, the function of *mirror* in determining the alula in *D. melanogaster*^13^ suggests that the alula may represent an evolutionarily reduced vannus. The ramifications of this are significant for reconstructing the history of the insect wing, because they suggest that the *D. melanogaster* wing blade is a lone remigium – only one-third of the archetypal insect wing **(Figure 1E)**. The dipteran jugum appears to have been lost entirely, except perhaps as a membranous haltere cover (calypter) in the Calyptratae clade, as speculated by Snodgrass.^15^ Thus, while flies have been an important model for characterizing genes and processes that build insect wings, a more comprehensive understanding of the development and evolution of insect wings requires work in species that have wings more representative of the complete ancestral blueprint.

## ACKNOWLEDGMENTS

We thank Kate Siegel, Nigel Williams, and Rick Fandino for assistance with rearing butterflies; Johanna Dela Cruz for help with confocal microscopy at Biotechnology Resource Center (BRC) Imaging Facility (RRID:SCR_021741) at the Cornell Institute of Biotechnology. We also thank Dr. Eirene Markenscoff-Papadimitriou for use the Leica Stellaris 5 confocal microscope for imaging. This work was supported by the United States National Science Foundation grants NSF IOS-1753559 and IOS-2128164 awarded to R.D.R., and a Cornell Summer Experience Grant and an Office of Undergraduate Biology fellowship to X.Y.

## MATERIALS AND METHODS

### Identity of *Iroquois* family gene *mirror* ortholog in *J. coenia*

We performed a reciprocal BLAST of amino acid sequences of *D. melanogaster Iroquois Complex* genes *araucan, caupolican*, and *mirror* to identify their orthologs in the latest *J. coenia* and *Heliconius erato lativitta* genome sequences on lepbase.org and *Tribolium castaneum* and *Apis mellifera* on NCBI. The amino acid sequences of the top hits – from *J. coenia* (JC_02265-RA, JC_02269-RA), *H. erato lativitta* (HEL_009655-RA, HEL_009656-RA), *A. mellifera* (LOC412840, LOC412839) and *T. castaneum* (LOC652944, LOC660345) were aligned to the *Iroquois* genes of *D. melanogaster* using MUSCLE.^17^ We used these alignments to build a maximum likelihood gene phylogeny on IQ-TREE 2^18^ using ModelFinder^19^ for estimating the most accurate substitution model for our amino acid sequences. We used FigTree to visualize the tree depicted in Figure S5.

### HCR fluorescent *in situ* hybridization of *mirror* mRNA

This protocol was adapted and modified from Bruce et al. (dx.doi.org/10.17504/protocols.io.bunznvf6). The coding sequence of JC_02269-RA was used by Molecular Instruments, Inc. (Los Angeles, CA, USA) to design a *mirror-*specific hybridization chain reaction (HCR) probe set (Lot #PRQ901). The probes are unique to the JC_02269-RA transcript as verified by BLAST search using the latest *J. coenia* genome assembly (v2) on lepbase.org, and do not share significant sequence similarity with any other transcripts, including the other *Iroquois* family member JC_02265-RA (see Figure S5). Last (5^th^) instar larval wing imaginal discs were dissected in cold 1x PBS. The dissected discs collected from left and ride sides of the larvae were randomly assigned to control or treatment batches. Both controls and treatment batches were fixed in 1x cold fix buffer (750uL PBS, 50mM EGTA, 250uL 37% formaldehyde) on ice. The discs were then washed three times in 1x PBS with 0.1% Tween 20 (PTw) on ice spending approximately 30sec – 2min for each wash. Tissues were then gradually dehydrated by washing in 33%, 66%, and 100% MeOH (in PTw) for 2-5min / wash on ice, and stored in 100% MeOH in -20°C for days until HCR protocol was started. On the day of the HCR protocol, the wing discs were progressively rehydrated in cold 75% MeOH, 50% MeOH, and 25% MeOH in PTw spending 2-5min per wash. After rehydrating, discs were washed once for 10min and twice for 5min with PTw. The discs were then permeabilized in 300-500μL of detergent solution (1.0% SDS, 0.5% Tween, 50mM Tris-HCL pH 7.5, 1mM EDTA pH 8.0, 150mM NaCl) for 30min at room temperature. While waiting, hybridization buffer (Molecular Instruments) was warmed to 37°C (200μL/tube). After permeabilizing with detergent solution, each tube of wing discs was incubated in 200μL of pre-warmed hybridization buffer for 30min at 37°C. Probe solution was prepared by adding 0.8pmol (0.8 μl of probe from 1uM stock solution) of each probe mixture to 200μL of probe hybridization buffer at 37°C. The pre-hybridization buffer was removed, and the probe solution was added. For negative control tissues, hybridization buffer without probes was added instead. All wing discs were incubated overnight (12-16hr) at 37°C. Before resuming the protocol, probe wash buffer (Molecular Instruments) was warmed to 37°C, amplification buffer (Molecular Instruments) was calibrated to room temperature, and a heat block was set to 95°C. The probe solution was removed and saved at -20°C to be reused. Wing discs were then washed four times in 1mL pre-warmed probe wash buffer at 37°C for 15min per wash. After the last wash step, discs were washed twice (5min/wash) with 1mL of 5x SSCT (5X sodium chloride sodium citrate, 0.1% Tween 20) at room temperature. The wing discs were pre-amplified with 1mL of equilibrated amplification buffer for 30min at room temperature. During this step, 2μL of each of hairpin h1 and 2uL of each hairpin h2 were mixed in 100μL of amplification buffer at 95°C for 90sec and cooled to room temperature in the dark for 30 min. For *mirror*-only *in situ* hybridizations, we used B1 hairpin amplifiers tagged with AlexaFluor 594 fluorophore (Molecular Instruments). For *mirror* and *wingless* double stains, we used *mirror* probes with B1 amplifier tagged with AlexaFluor 647 fluorophore and *wingless* probes (Lot #PRG129) with B3 hairpin amplifiers tagged with AlexaFluor 546 fluorophore. After 30min, the pre-amplification buffer was removed from the discs, hairpin solution added, and the tissues were then incubated in the dark at room temperature for 2-16hr. The hairpins were removed and saved at -20°C to be reused later. Excess hairpins were removed by washing 5 times (twice for 5min, twice for 30min and once for 5min) with 1mL of 5x SSCT at room temperature. The wing discs were then incubated in 50% glycerol solution (in 1X PBS) with DAPI (0.01μg/mL) overnight at 4°C. The wing discs were then mounted and visualized on a Zeiss 710 or Leica Stellaris 5 confocal microscope.

### CRISPR-Cas9 mediated mutagenesis of *mirror* in *J. coenia*

Two single-guide RNAs (sgRNAs; sgRNA1-5’-GAATGGACTTGAACGGGGCA; sgRNA2-5’-AGAAACAGGGTCGATGATGA) targeting the homeobox domain of *mirror* were mixed with 500ng/μl of Cas9 nuclease and injected to *J. coenia* eggs 0.5-4hr after oviposition (n = 1042 eggs) as previously described. ^20^ Injected G0 individuals were reared on standard artificial diet, until they emerged and were immediately frozen in -20^°^C upon emergence. Wings of surviving G0 adults (n=99) were assayed for anomalies to detect mosaic knockout (mKO) phenotypes (n=29). Injection results and mutation phenotype frequencies are detailed in Table S1.

### Validating *mirror* mutants by genotyping

DNA was extracted using E.Z.N.A. Tissue DNA Kit (Omega Bio-Tek) from the thorax of individuals that showed mutations in the wings. Extracted genomic DNA was amplified using a pair of primers flanking the sgRNA cut sites (Forward: 5’-CGCTTGTGCCCACCTTAAAC, Reverse: 5’-GTATGGCTCGGGGGATTCTG). Amplified DNA was run on a 2% agarose gel and were excised and purified using the MicroElute Cycle-Pure Kit (Omega Bio-Tek). Purified DNA was Sanger sequenced by Cornell Institute of Biotechnology. Example mutant alleles of *mirror* are shown in Figure S6.

## AUTHOR CONTRIBUTIONS

Conceptualization, M.C and R.D.R; Methodology, M.C and R.D.R; Investigation, X.Y.Y, M.C, N.K.B , G.C.H and C.A.A; Writing-Original Draft, M.C, X.Y.Y and R.D.R; Writing-Review & Editing, M.C and R.D.R; Visualization, M.C and X.Y.Y; Funding Acquisition, R.D.R and X.Y.Y; Supervision, M.C and R.D.R

## SUPPLEMENTAL INFORMATION

**Figure S1.**
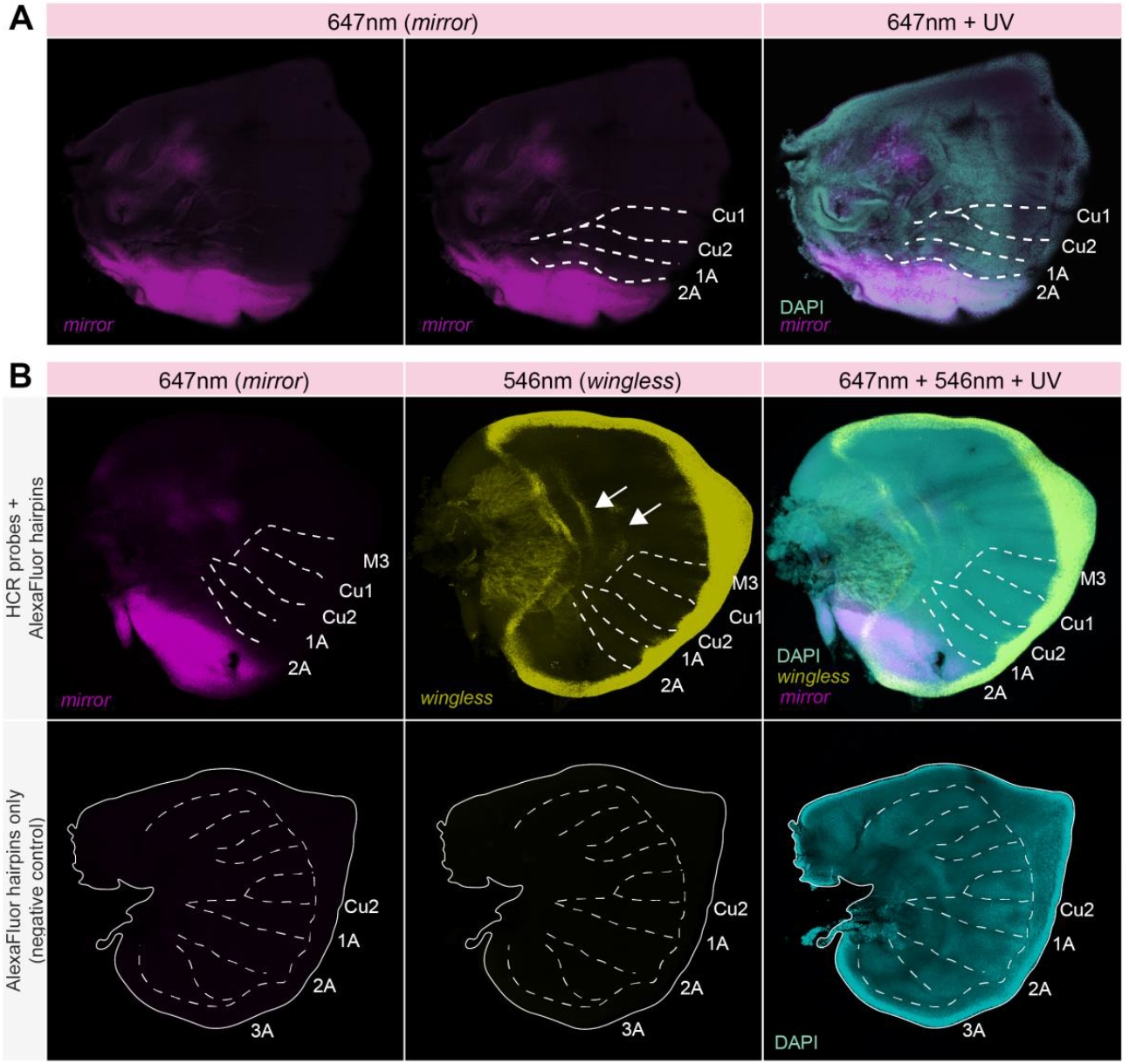
HCR *in situ* hybridization of *mirror* highlight posterior domain expression in *J. coenia* wing imaginal discs. Related to Figure 2A. **(A)** *mirror* is expressed posterior to the 2A vein in last-instar forewing imaginal discs of *J. coenia*. **(B)** Double *in situ* of *mirror* (Channel 1: 647nm) and *wingless* (Channel 2: 546 nm) in last instar *J. coenia* hindwing imaginal discs. Expression of *wingless* observed by HCR precisely matches expression in wing margin peripheral tissue and discal bands (white arrow) as previously described from chromogenic in situs^5^, thus providing a positive control for our HCR protocol. Bottom panels depict negative controls, lacking HCR probes, that show no signal in the 647nm or 546nm channels. Dashed lines show position of selected veins.

**Figure S2.**
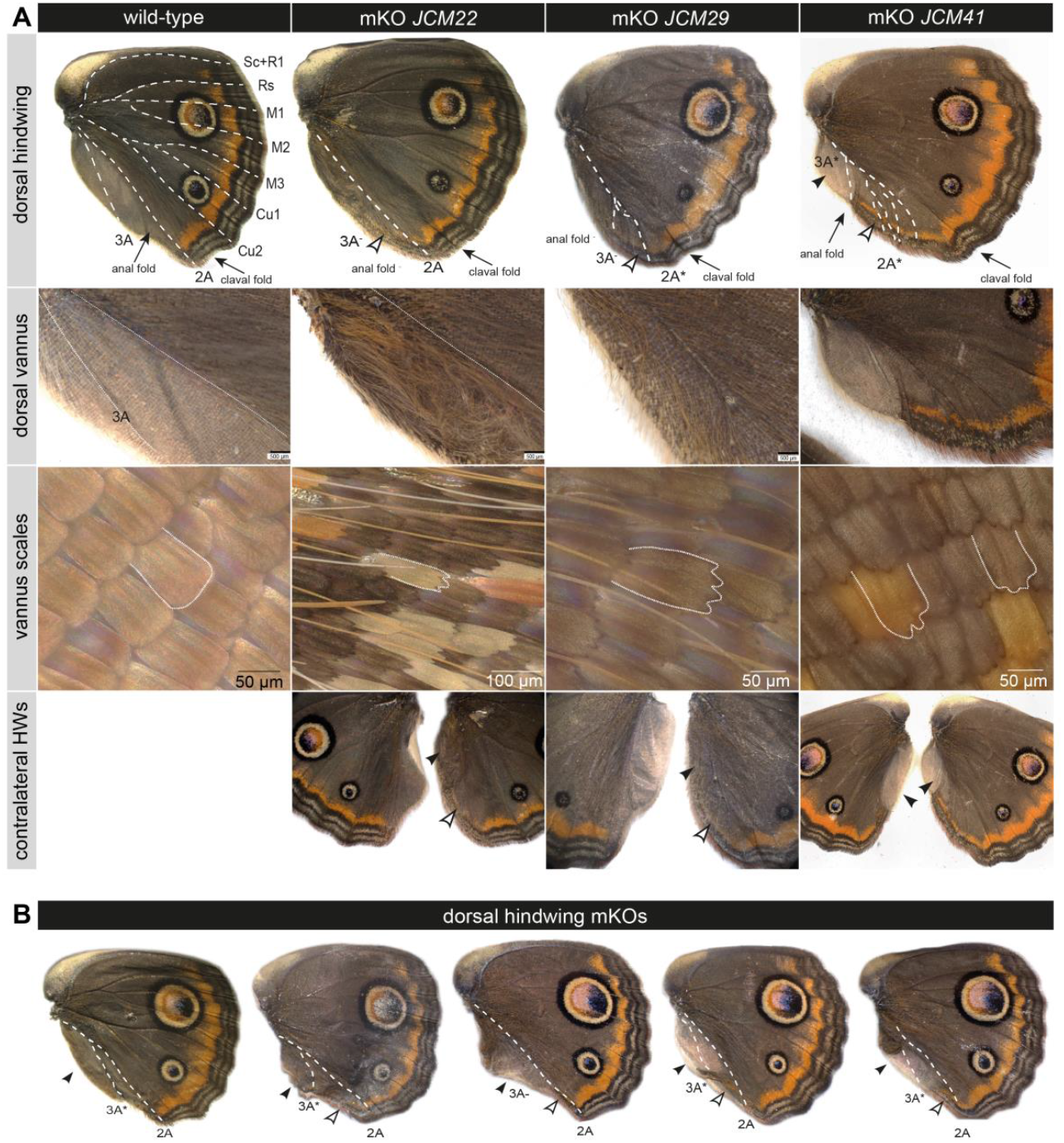
mKOs of *mirror* result in partial or entire loss of posterior wing domain identity in dorsal *J. coenia* hindwings. Related to Figure 2B-E. **(A)** Comparison of wild-type dorsal hindwing to mutant wings reveals complete or partial transformation of the vannus (black arrows), expansion of color patterns into the posterior wing margin (white arrows) along with vein aberrations (denoted by *), or total loss (denoted by -). The middle panels show magnified images of the posterior wing margin and vannus, with scale morphology details, in wild-type and mKO mutants (complete transformation in *JCM22* and *JCM29*; partial transformation in *JCM41*). The bottom panels show contralateral hindwing pairs from the same individuals showing asymmetry in clonal mutations affecting posterior wing margin phenotype. **(B)** Additional *mirror* mKO hindwings show vannus transformation phenotypes, annotated as described above.

**Figure S3.**
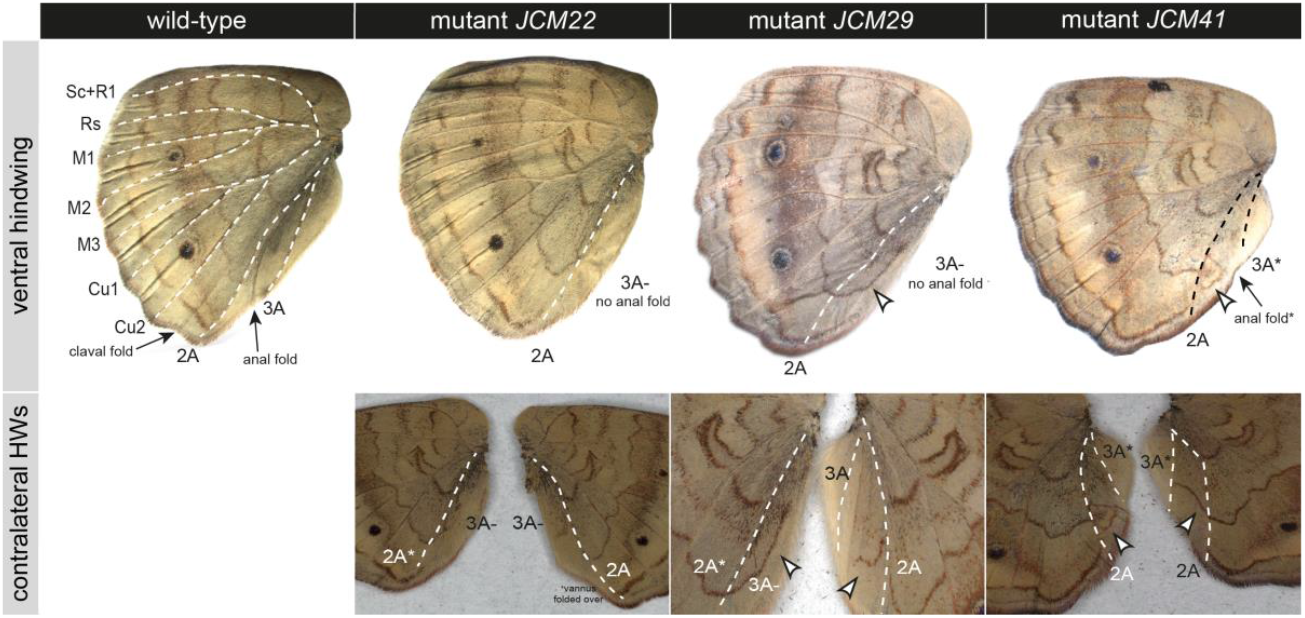
Related to Figure 2B-E. mKOs of *mirror* result in partial or entire transformation of posterior wing domain. *mirror* mKOs in ventral *J. coenia* hindwings resulting in the expansion of color pattern elements (white arrows) as seen in asymmetric contralateral pairs from the same mutant individuals. Abnormal venation annotated by (*), missing venation annotated by (-).

**Figure S4.**
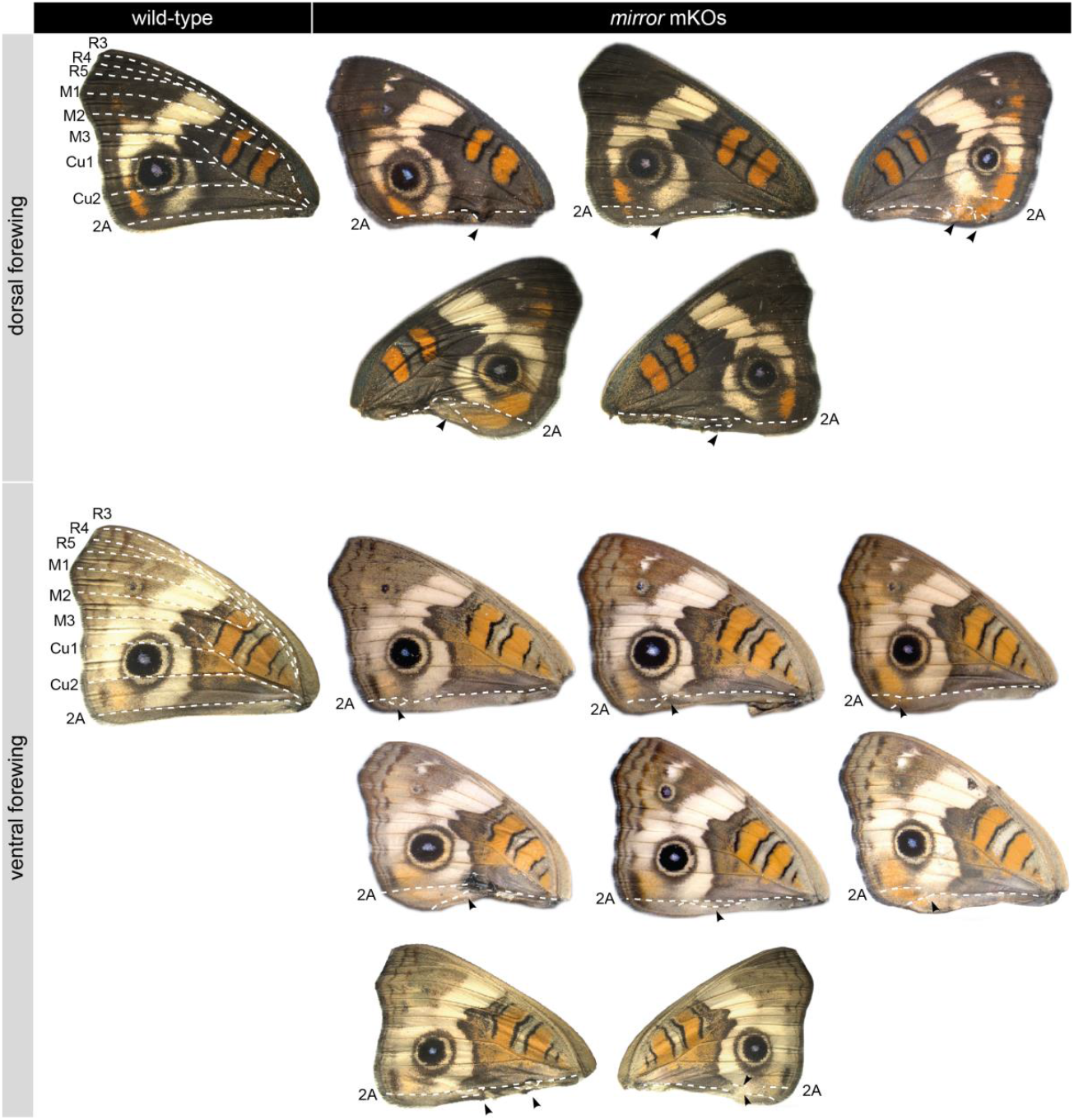
*mirror* mKOs show posterior margin anomalies *J. coenia* forewings. mKOs of *mirror* result in anomalies along the forewing 2A vein, including vein bifurcation and ectopic vein fragments, which are often associated with color pattern disruption. Dashed lines show the position of 2A. Black arrows highlight 2A abnormalities.

**Figure S5.**
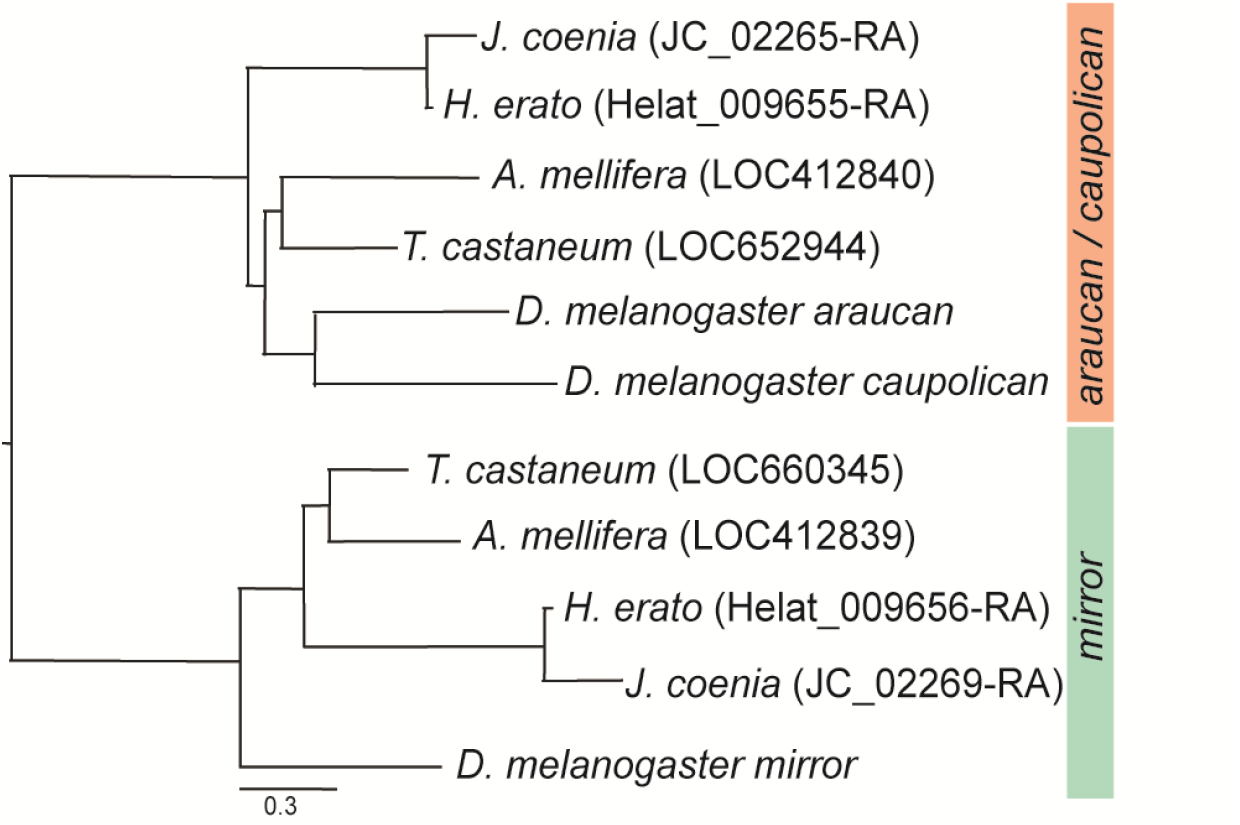
Identification of *mirror* ortholog in *J. coenia*. A maximum likelihood phylogeny of *J. coenia, H. erato lativitta, Tribolium castaneum, Apis mellifera*, and *D. melanogaster Iroquois Complex* genes confirms that JC_02269-RA is the ortholog of *mirror*.

**Figure S6.**
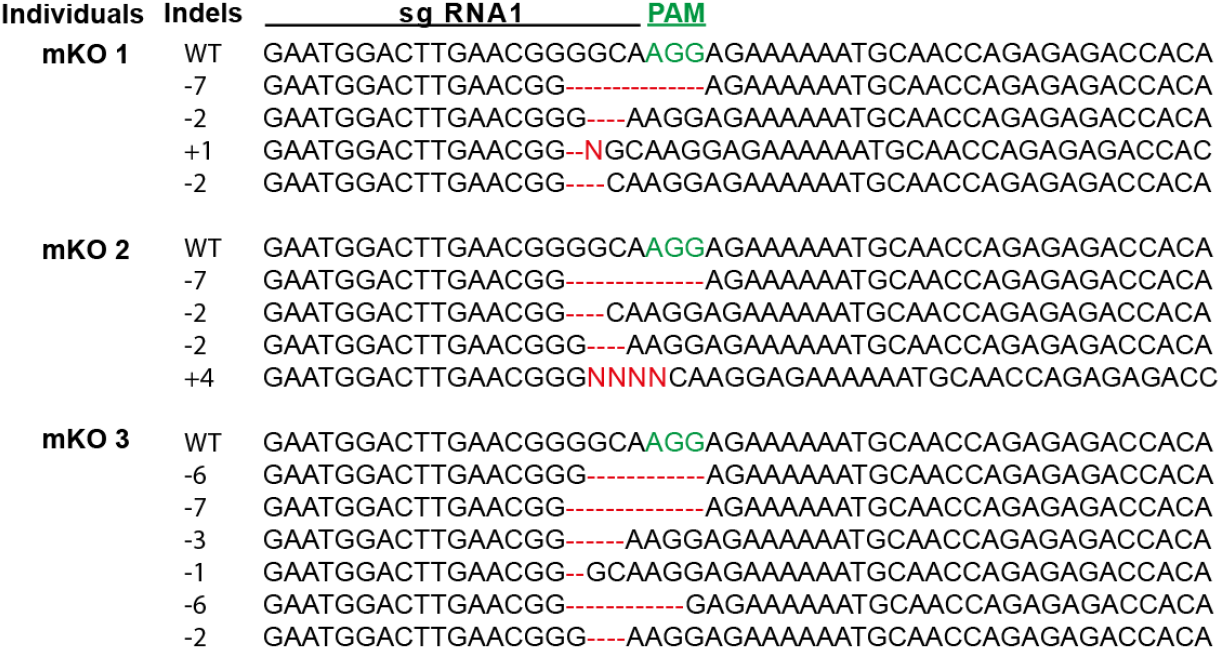
Sanger sequencing confirms mutations at CRISPR sgRNA site in *mirror* mKOs. Sanger sequencing results from three different *mirror* mutant individuals. Each line denotes a different wild-type (WT) or mutant allele with indels indicated in red, and the PAM site in green.

**Table S1.** *mirror* CRISPR/Cas9 injection results.

| **sgRNA**<br>**concentration**<br>**(ng/μL)** | **Eggs**<br>**Injected** | **Hatched** | **Adults**<br>**Emerged** | **Mutants** |
| --- | --- | --- | --- | --- |
| 487 ng/μL | 142 | 25 | 4 | 2 |
| 391 ng/μL | 96 | 33 | 6 | 3 |
| 335 ng/μL | 163 | 32 | 15 | 4 |
| 281 ng/μL | 98 | 11 | 1 | 0 |
| 175 ng/μL | 149 | 52 | 19 | 8 |
| 125 ng/μL | 252 | 79 | 24 | 9 |
| 88.5 ng/μL | 142 | 41 | 30 | 3 |
| **Total** | **1042** | **273** | **99** | **29** |

